# Reciprocal social ties drive stability within a social network

**DOI:** 10.1101/2020.11.06.371567

**Authors:** Roslyn Dakin, Paisley Clunis, Emily M. Cornthwaite, T. Brandt Ryder

**Affiliations:** Department of Biology Carleton University 1125, Colonel By Drive, Ottawa, Ontario, Canada K1S 5B6; Migratory Bird Center, Smithsonian Conservation Biology Institute National Zoological Park, Washington, DC, USA 20013; Bird Conservancy of the Rockies Fort Collins, Colorado, USA 80525

**Keywords:** dynamic network, directed network, manakin, social connectivity, reciprocity, bidirectional, sociality

## Abstract

Social reciprocity is thought to be the most important driver of cooperation among non-kin, but its effects within complex social networks where individuals choose their partners have not been studied. Here, we test whether reciprocation can explain social dynamics in a bird species where males form coalitions to court females, the wire-tailed manakin (*Pipra filicauda*). In our study population, approximately a third of male-male social interactions involve one male leaving his territory to interact and display with the owner of another territory; this directional behaviour provides benefits to the recipient at a cost to the visiting male. We characterize social ties among pairs of territory holders according to the directionality of their interactions, and show here that territory holding males engaged in far more reciprocated (bidirectional) partnerships with other territory holders than expected by chance. Reciprocated partnerships were also stronger (i.e., the partners interacted more frequently) than non-reciprocated partnerships, controlling for spatial proximity of the territories. An individual-level analysis revealed that a male’s social contribution to a given partner was predictive of the number of social interactions he received from that same partner. Finally, we show that reciprocation predicted the long-term stability of social partnerships one year later. Together, these results demonstrate that reciprocity is associated with stability in complex social networks where partner choice is flexible.

**LAY SUMMARY:** *When animals are free to choose their partners, how does reciprocity influence social dynamics?:* In the wire-tailed manakin, males form coalitions with other males to help attract mates. We tracked male social networks and found that some males form reciprocal partnerships where each male leaves its territory to display at the other’s territory. These bidirectional partnerships are particularly strong and stable, indicating that reciprocity is associated with stability in complex social networks.

## Introduction

Helping behavior among non-kin is found in many taxa, including mammals, birds, fish, and invertebrates (Hart and Hart 1992; McDonald and Potts 1994; Griffiths and Magurran 1997; Backwell et al. 1998; DuVal 2007; Rutte and Taborsky 2007; Brucks and von Bayern 2020; Moore et al. 2020). One of the primary mechanisms thought to drive the evolution of cooperative behaviour in non-kin is social reciprocity (Trivers 1971; Roberts and Sherratt 1998; Taborsky et al. 2016). Recently, several pioneering studies have used careful experiments to demonstrate that animals can both detect and respond to help from unrelated social partners (Rutte and Taborsky 2007; Cheney et al. 2010; Brucks and von Bayern 2020; Carter et al. 2020; Engelhardt and Taborsky 2022). In nature, social relationships often occur in the context of complex social networks, where individuals are free to choose their partners, and where social structure changes dynamically through time (Hart and Hart 1992; Croft et al. 2008; Fehl et al. 2011; Ilany et al. 2015). In natural social systems, an individual’s social investment into one partner (or social tie) often occurs at the cost to other partnerships and to self-maintenance behaviour (e.g., foraging, reproduction). Consequently, social reciprocity is predicted to drive long-term patterns of group-level social structure and stability (Dakin and Ryder 2020; Yamamoto and Ishibashi 2022). Despite the well-known importance of reciprocity in evolutionary and ecological theory (Fehl et al. 2011), previous studies have not yet examined how mutual social investment affects dynamics within a complex social network.

Here, we test the hypothesis that mutual social investment can explain social dynamics in a complex social system of wire-tailed manakins *Pipra filicauda* (Ryder et al. 2008). This species is part of a large clade of manakins characterized by elaborate male displays that are considered to be some of the most complex of all passerine birds (Schwartz and Snow 1978; Prum 1990; Heindl 2002; Ryder et al. 2008). In the wire-tailed manakin, males form cooperative partnerships with non-kin and perform coordinated displays at specific display perches within their lek territories. A typical male will have several social partners that he displays with repeatedly, within and across breeding seasons (Ryder et al. 2008; Ryder et al. 2011; Dakin and Ryder 2020). Previous studies have shown that an individual male’s social centrality within the broader male coalition network is a strong predictor of his ability to acquire a territory and achieve long-term siring success (Ryder et al. 2008; Ryder et al. 2009). Male-male partnerships in this system are also temporally stable across multiple years (Ryder et al. 2011; Dakin and Ryder 2020), as observed in other manakins (Trainer 2002).

In the wire-tailed manakin, there are two male status classes: the dominant males who hold territories, and the subordinate floater males who do not yet hold a territory (Heindl 2002; Ryder et al. 2008; Ryder et al. 2009; Ryder et al. 2011). Although most social interactions occur between a floater and a territory-holder, approximately a third of social interactions involve one male leaving his territory to interact and display with the owner of another territory in a directed interaction (Dakin and Ryder 2020). Importantly, the visiting male in this scenario takes on a crucial cost: by leaving his own territory, he foregoes potential mating opportunities and beneficial social ties that could have occurred there (Ryder et al. 2008; Ryder et al. 2009). On the other hand, the recipient male in this scenario is expected to gain a fitness benefit, because social displays on a male’s own territory are associated with increased siring success (Ryder et al. 2008; Ryder et al. 2009). These directed interactions provide an opportunity to investigate how often reciprocity occurs, and to test the prediction that reciprocity is associated with stability in social network dynamics (Dakin and Ryder 2020).

We categorized social partnerships among territory-holding wire-tailed manakins as either mutually reciprocal or one-directional (Fig. 1A). A mutually reciprocal partnership is a bidirectional relationship in which the two territory-holders interacted with each other mutually on their two respective territories. One-directional partnerships are those in which only one of the territorial males visited the other male’s territory to interact socially with him, and the reverse did not occur. We determined the directionality of manakin partnerships over both weekly and annual timescales. Our analysis here evaluates the following four questions: (1) Do reciprocated cooperative partnerships occur more often than expected by chance, based on individual spatiotemporal behavior? (2) Are reciprocated partnerships preferred, as defined by having a higher social interaction frequency, as compared to one-directional partnerships? (3) At an individual level, does the magnitude of one male’s social contribution to a given partner predict the contribution he received from that partner? (4) Are reciprocated partnerships more temporally stable in the long-term than one-directional partnerships? Together, these analyses allow us to test whether reciprocity is a mechanism of social preference and temporal stability within animal social networks.

**Figure 1.**
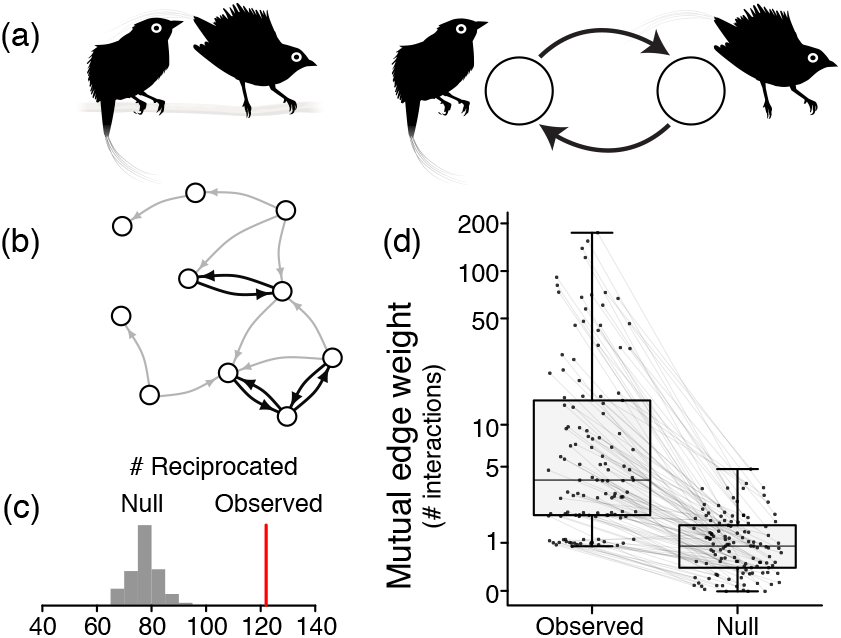
Social relationships can be one-directional or reciprocated. (A) Male manakins form cooperative partnerships with other males to perform coordinated courtship displays. A reciprocated partnership occurs when two territory holders interact with each other mutually on their two respective territories. In the network diagrams, the circles represent each male’s territory, and the arrows represent directional social interactions that originate from the visitor and point toward the recipient. (B) Example of a directed social network among 10 territory-holding males. (C) In total, we observed 122 reciprocated annual partnerships (indicated in red), which is 56% more than expected based on a null permutation. (D) The mutual edge weights of reciprocated partnerships were also far greater than expected by chance. The boxplots show mutual edge weights with lines connecting the observed value for each partnership with its null expectation (n = 122 reciprocated annual partnerships). See also Table S1.

## METHODS

### (a) Field work

We studied a population of wire-tailed manakins at the Tiputini Biodiversity Station in Orellana Province, Ecuador (0° 38’ S, 76° 08’ W, ∼20m elevation). Field work took place over three breeding seasons (December to March) from 2015 to 2018. At the beginning of each field season, wire-tailed manakins were captured using mist nets and tagged with a unique combination of color bands as well as a coded nanotag (NTQB-2, Lotek Wireless; 0.35 g). Nanotags transmitted a unique VHF radio signal throughout the field season. We sampled males from 11 leks at the field site, with lek sizes ranging from 4-14 territories (and typically, the number of males using a lek is about 2x the number of territories). To record male social interactions on lek territories, we placed proximity data-loggers (SRX-DL800, Lotek Wireless) within each territory on a lek for a recording session of ∼6 consecutive days (± SD 1 day; loggers were active between 6:00 to 16:00 each day). Each lek was monitored for 3–4 recording sessions per field season, distributed approximately evenly throughout the reproductive period from December to March. Territory ownership was determined through direct observation and confirmed in the proximity data. To determine the distance between territories, GPS latitude and longitude coordinates were recorded at a location near the display perch within each territory. In total, we monitored between 48–54 territory-holders per year (see Table S1).

### (b) Social data

We used an algorithm described in (Dakin and Ryder 2018) to identify social interactions between two male manakins within a display territory. The algorithm defined a social interaction to have occurred when two males pinged a data-logger within a threshold of 45 s and 10 signal strength units, which indicates spatial proximity of two males < 5 m of each other near the display perch (Dakin and Ryder 2018; Dakin et al. 2021). This method has been shown to reliably detect social interactions between male wire-tailed manakins in ground-truthing experiments (Ryder et al. 2012; Dakin and Ryder 2018). All data processing and analyses were performed in R 3.6.3 (R Core Team 2020). Here, we focus on interactions of interest between pairs of territory holding males. We define a directed social interaction among territory holders as occurring when two territory-holders engaged in a social interaction on one of their respective territories (Pinter-Wollman et al. 2014). A partnership between two territory-holders was defined as one-directional if all of their social interactions during a given time period occurred on only one of their respective territories. A partnership was defined reciprocated (or bidirectional) if the two males interacted on both of their territories. We computed both directed and undirected edge weights based on the frequencies of these social interactions. The undirected edge weight was the total frequency of social interactions for a dyad within a given time period (either weekly or annual); the directed edge weights refer to the frequencies occurring on each recipient male’s territory. An important indicator of directional symmetry for reciprocated partnerships is the minimum edge weight that occurred in both directions. Hence, we defined an additional metric that we refer to as the “mutual edge weight”, defined as the smaller of the two directed edge weights for a given dyad. Recent experiments have established that spatial proximity is an important driver of reciprocal relationships (Razik et al. 2022). To consider spatial distances as a potential determinant of behavior, we estimated the Euclidean spatial distance between each pair of territories from their latitude and longitude coordinates using the sp package 1.4-2 (Pebesma and Roger 2020) in R.

### (c) Raw data stream permutation

We compared the occurrence of reciprocated partnerships with the number expected to occur by chance using a null model of territory visitation (Farine 2017; Dakin et al. 2021). The null model was based on a permutation of the territory visits performed by each male, as described in (Dakin et al. 2021). Note that we use “territory visit” here to refer to the detection of an individual male on *any* territory, independent of social context. The null permutation preserved each male’s spatial and temporal distribution of territory visits, including the dates of each visit, and the visit durations, but randomly sampled the visit start times from the distribution of actual start times observed in the data (Dakin et al. 2021). Hence, the null model preserved each male’s spatiotemporal behavior (i.e., the same territories were visited, the same number of times, and for the same duration), but ensured that the exact timing of visits was independent of the presence of other males. This approach previously demonstrated that manakin social partnerships have much greater undirected edge weights than expected (Dakin et al. 2021). For the present study, we used 100 permuted null datasets to determine the expected number and directionality of annual partnerships among territory holders, and their edge weights (undirected and mutual; based on frequency of social interactions, as defined above). We then compared these expected values from the permutation analysis with the actual values observed in the manakin data.

### (d) Predictors of whether a partnership was reciprocated

We used multilevel models at the dyad level to determine the factors that could predict whether or not a partnership was reciprocated within a particular recording session. The response variable for this analysis was the partnership type (reciprocated vs. one-directional), modelled in a binomial analysis in the lme4 package 1.1-23 (Bates et al. 2020) in R. Because this analysis focused on a dyad-level property, the random effect accounted for the dyad identity, based on the combined identities of the two participating males. To examine the factors that predict reciprocation, we examined two fixed effect predictors: (i) the undirected edge weight (log-transformed) and (ii) the spatial distance between the two males’ territories. We also accounted for fixed effects of field season (three levels, one for each year) and sampling effort, as defined by the total number of hours that the two territories were recorded. To ensure that this analysis only considered partnerships that could have been reciprocated, we limited it to the 377 session-partnerships that had an edge weight ≥2 (the minimum required for reciprocation), and where both males’ territories were monitored throughout the season.

### (e) Magnitude of individual contributions in reciprocated partnerships

The goal of this individual-level analysis was to describe the factors that predict a given territory-holder’s social contribution to his territorial partners within a recording session. Hence, we use ‘strength’ here to refer to the social interaction frequency of a particular individual (i.e., the sum of the edge weights for that individual’s node in the social network). This analysis was run using a multilevel model in the lme4 package in R, with the response variable as the relative amount that a focal male contributed to a specific partner. To calculate this variable (relative amount contributed), we took the focal male’s social interactions contributed to partner *x_i_*, divided by the focal male’s total social interactions away from his own territory. Note that “contributed” here refers to times when the focal male went to the partner’s territory and interacted with him there. The main predictor of interest was the relative amount that the focal male received on his territory from partner *x_i_* (i.e., strength received from *x_i_* expressed relative to the focal male’s total social interactions received). Thus, this analysis tested whether the focal male’s social contribution to a particular partner was correlated with the amount received (reciprocal social investment) from that partner. Additional fixed effects accounted for field season (three levels, one for each year), the spatial distance between territories, and sampling effort. We also included the duration of visits by floater males (i.e., males that do not hold territories) to the focal male, and the duration of visits by floater males to partner *x_i_* as two additional fixed effect predictors (both expressed as the natural log of the total number of floater pings). This allowed us to account for the possibility that social contributions among territory holders may depend on the activity of these other male social partners within the lek. The identities of the focal and partner male were included as crossed random effects. We limited this analysis to reciprocated session-partnerships, represented in a sample size of 326 contributions within 163 partnerships among 72 males.

### (f) Predictors of long-term partnership stability

To evaluate whether reciprocity leads to long-term stability of male-male partnerships, we analyzed the interaction frequency of a partnership in the following year, after an initial designation of partnership type. The response variable was the partnership’s undirected edge weight in the second year, *t + 1*, analyzed in a Gaussian model in lme4. To meet the assumptions of the Gaussian model, the response variable was log-transformed. To test whether previous reciprocation could predict long-term dynamics, the model included a fixed effect of partnership directionality in the initial year *t* (either reciprocated or one-directional). The other fixed effects in the model included field season, the spatial distance between territories, and the edge weight of the partnership in the initial year *t* (log-transformed). Hence, this analysis tested whether edge weight in year *t + 1* was associated with previous reciprocation in year *t*, after accounting for the previous edge weight. Because this analysis focused on a dyad-level property, the random effect accounted for identity of the dyad as describe above. The criteria for inclusion in this analysis was that the year *t* edge weight had to be ≥2 (i.e., the partnership had to have occurred often enough that it could have been reciprocated in year *t*), and that both males had to be tagged and monitored in the subsequent field season; the resulting sample size was 109 annual-partnerships.

## RESULTS

### (a) Occurrence of reciprocated partnerships

We recorded over 9,500 directed social interactions between two territorial male wire-tailed manakins and identified 122 bidirectional annual-partnerships that were reciprocated at least once. The number of reciprocated partnerships was 56% more than expected, based on a null model (Fig. 1c). By contrast, we observed far fewer one-directional partnerships among territory holders than expected by chance (Table S1). This indicates that reciprocity is not occurring at random; rather, it is a common feature of the wire-tailed manakin social system. The average territory holder had about 4 (± 2.0 SD) territorial partners per year, 2 (± 1.4 SD) of which were reciprocated, and 2 (± 1.6 SD) of which were one-directional.

The mutual edge weights of reciprocated partnerships were typically 6-to-14-fold greater than expected in the null model, demonstrating that these social partnerships are far more symmetrical than expected (Fig. 1d). For example, 58 partnerships had an annual mutual edge weight ≥ 5, a level of symmetry that was never observed in the null model (Fig. 1d). Moreover, we found that reciprocated social partners interacted 10-to-20-fold more than expected by chance (based on undirected edge weights; see Table S1). One-directional social partners also interacted more than expected, but to a much lesser extent, with only a 3-to-7-fold difference as compared to the expected values (see Table S1).

### (b) Predictors of whether a partnership was reciprocated

Reciprocation was more likely to occur within partnerships that interacted frequently, as compared to partners that interacted rarely (Fig. 2). This strong association between reciprocation and edge weight also holds while accounting for the spatial proximity of the two territories, which was another significant predictor of reciprocation. Pairs with closer territories were more often reciprocal (Table S2).

**Figure 2.**
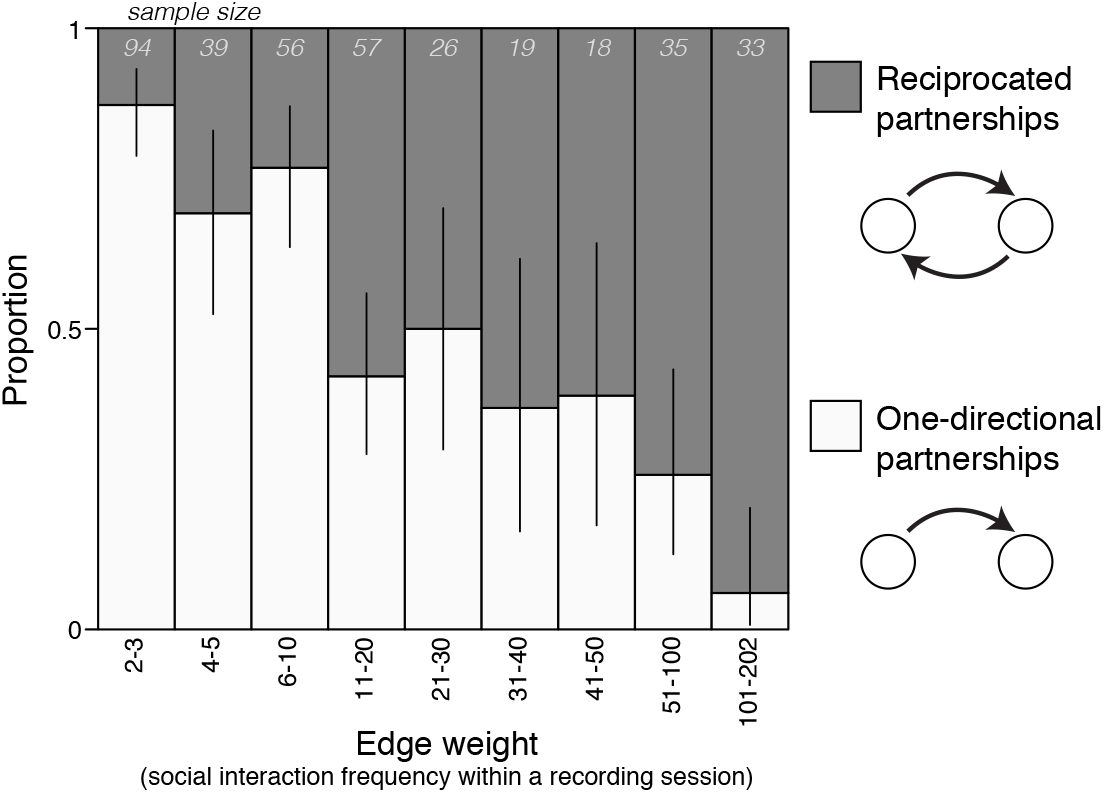
Reciprocated dyads interact more frequently. Reciprocation was more likely to occur among partners that interacted more frequently with each other (greater undirected edge weight). The positive association between reciprocation and edge weight holds while accounting for the spatial proximity of the two males’ territories, field season, recording duration, and the identities of the two partner males. The number of session partnerships in each column is provided along the top of the graph (total = 377 session partnerships with an edge weight >= 2, involving 80 males). Error bars show the 95% confidence intervals for the proportion estimated in each column. Note that while edge weight is binned here for visualization purposes, it was treated as a continuous variable in the statistical analysis (see Table S2).

### (c) Magnitude of individual contributions in reciprocated partnerships

There were two main factors that predicted a male’s social contribution to a given partner. The first predictor was the relative social contribution received from that partner: focal males directed more of their social interactions toward partners they had received more from (Fig. 3; Table S3). The second major predictor was the spatial proximity of their territories: males contributed more to closer neighbors (Table S3). The magnitude of social contribution was not significantly associated with the frequency of floater male occurrence at either territory (Table S3).

**Figure 3.**
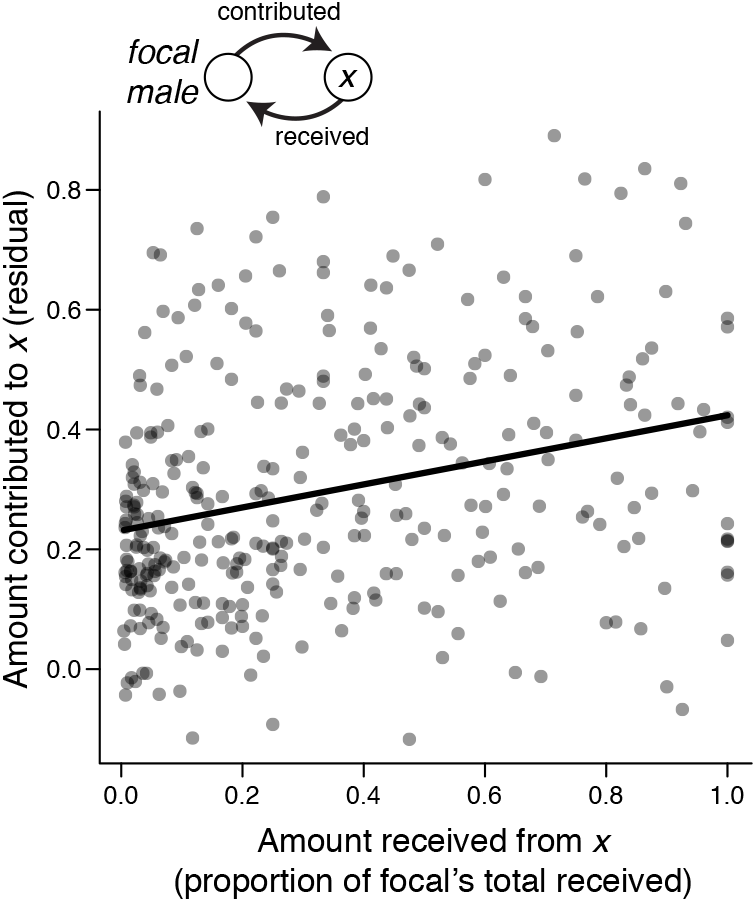
A male’s contribution to a given social partner is correlated with the amount he received. Following the diagram on the top left, this analysis evaluated whether a focal male’s contribution to partner *x* was predicted by the relative number of social visits received from partner *x*. The y-axis shows partial residuals for a male’s social contribution while controlling for other factors in the analysis, including proximity, field season, recording duration, floater male visits, and the identities of the two partner males (see Table S3 for details). Note that this analysis is limited to session partnerships that were reciprocated (n = 326 individual contributions within 163 session partnerships, among 72 males).

### (d) Predictors of long-term partnership stability

Partnerships that were reciprocated in year *t* maintained far greater edge weight (interaction frequency) the following year than did one-directional partnerships (Fig. 4). This result held even when accounting for territory proximity and the previous year’s edge weight, both of which were also positive predictors of edge weight in the following year (Table S4).

**Figure 4.**
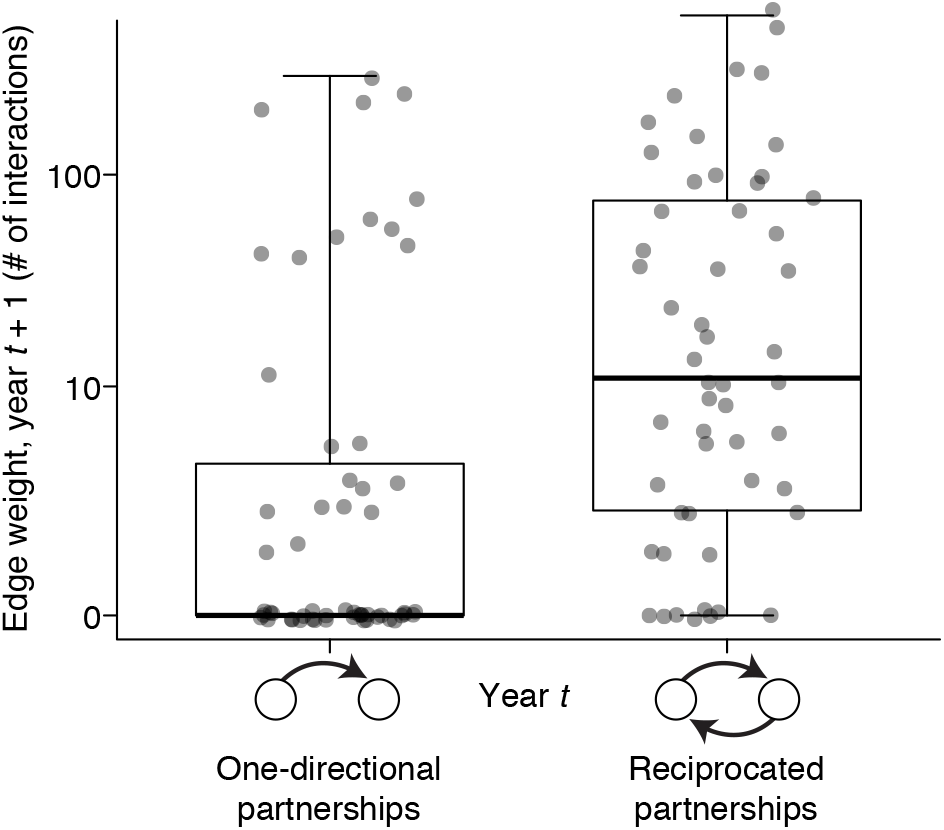
Reciprocated dyads maintain greater stability through time. Social dyads that were reciprocated in year *t* had significantly stronger edge weights (interaction frequencies) in year *t* + 1 (n = 109 annual partnerships among 38 males). This positive association holds while accounting for the undirected edge weight in year *t*, spatial proximity of the two males’ territories, field season, and the identities of the two partner males (see Table S4). Note that this analysis is limited to dyads that interacted at least twice in year *t* and that were both tagged and monitored over the following year.

## DISCUSSION

Reciprocation has been proposed to drive the evolution of cooperation and determine the structure and dynamics of partner choice within cooperative social networks (Fehl et al. 2011; Dakin and Ryder 2020). We used an analysis of directed social ties to test how reciprocation predicts social network dynamics in a wild population of cooperative birds, where individuals were free to choose their social partners. We found that the social networks of territory holding male manakins contained many more reciprocated (bidirectional) partnerships than expected, and fewer one-directional partnerships than predicted by a null model of individual spatiotemporal behaviour. Moreover, mutual edge weight, a key measure of social symmetry, was also found to be much higher than expected. Our analysis demonstrates that there are two main predictors of whether or not a social interaction was reciprocated: the overall frequency of social interactions between the two partners, and the spatial proximity of their territories. Male-male pairs that were located physically closer to each other, and those with a higher frequency of social interaction, were more likely to have a reciprocated relationship. Together, these findings indicate that reciprocation is either a cause of, or a marker of, social preference. They also indicate that spatial structure can also influence the costs and benefits of an individual’s social choices (Backwell et al. 1998; Webber and Vander Wal 2018; Bonar et al. 2020).

Previous studies of wire-tailed manakins have shown that a male’s social connectivity on his own territory is correlated with reproductive success (Ryder et al. 2008; Ryder et al. 2009). Here, we show that a male manakin’s social contribution to another territory-holding partner was positively correlated with the number of social interactions he received from that partner, independent of spatial proximity. Collectively, these findings suggest that males who form strong social bonds with other territory holders may receive delayed benefits through reciprocated social interactions that “return the favour”. This result adds to the growing literature that animals can respond in-kind to others’ social contributions (Rutte and Taborsky 2007; Cheney et al. 2010; Brucks and von Bayern 2020; Carter et al. 2020; Mishra et al. 2020; Lalot et al. 2021). Importantly, the present findings demonstrate that reciprocation can drive social investment in natural systems where individuals allocate their time towards multiple partnerships within a complex network.

How does reciprocity influence the long-term dynamics of social ties? We found that reciprocated cooperative partnerships retained greater social interaction frequencies one year later, even after accounting for positive effects of the initial social interaction frequency and territory spatial proximity. This finding indicates that cooperative relationships that are reciprocal are more likely to be maintained at a high rate in the future. Although we lack the ability to experimentally manipulate social bonds, this longitudinal analysis is consistent with two non-exclusive hypotheses, as mentioned above: (1) reciprocation may directly enhance social preference for particular partners, and (2) reciprocation may be a marker of (or consequence of) more general social compatibility between two individuals. Under the first hypothesis, reciprocation in the past is a direct cause of future social preference (Roberts and Sherratt 1998). The fact that reciprocation predicted future dynamics after accounting for the previous social interaction frequency is suggestive of this first possibility, and is consistent with recent controlled experiments in other species (Carter et al. 2020). An additional possibility is that reciprocation in year *t* may be a consequence of more general social compatibility (Busia et al. 2017; Ellis et al. 2019), which may in turn drive the enhanced stability of particular social bonds. Although we are unable to experimentally disentangle these two possibilities with the present data, future studies may be able to address this by the comparison of data on the fine-scale temporal dynamics of natural systems with carefully constructed simulation models.

Our results raise further questions about the behavioral mechanisms underlying reciprocation in the wild. For example, the results of our individual-level analysis suggest that wire-tailed manakins may recognize and distinguish the quality of cooperative social partnerships with particular individuals (Cheney et al. 2010). This raises the question of whether individuals within complex social networks can allocate their social contributions strategically (Ellis et al. 2019). Although not a main focus of our study, our results also emphasize the importance of physical proximity in determining the directionality and dynamics of social network ties (Webber and Vander Wal 2018; Webber et al. 2023). Because a positive correlation exists between proximity and the probability of an initial interaction occurring within a pair, spatially constrained social systems may have a greater potential for reciprocated interactions.

Historically, research on the ecology of cooperation among non-kin has focused on distinguishing reciprocal altruism from other mechanisms, such as by-product mutualism (Nowak 2006). Recent studies highlight how reciprocation can often be graded and dynamic in time (Roberts and Sherratt 1998; Carter et al. 2020). We show here that reciprocation occurs often in a complex social network, that it is either a cause of or a marker of social preference, and that it is associated with long-term social stability of cooperative partnerships (Fehl et al. 2011; Dakin and Ryder 2020). More broadly, social network structures are an important framework for understanding the movement ecology and behaviour in the wild. We propose that reciprocal network ties may often be stronger and more stable, and that the emergent properties of social networks will often depend on the composition and symmetry of these bidirectional ties.

## ETHICS STATEMENT

All research was approved by the Smithsonian ACUC (protocols #12-23, 14-25, and 17-11) and the Ecuadorean Ministry of Environment (MAE-DNB-CM-2015-0008).

## DATA ACCESSIBILITY

All data and R scripts are available at: https://figshare.com/s/f225fefb1ec4906e0353 The repository will be made public when the final version of the study is published.

## AUTHORS’ CONTRIBUTION

All authors designed the study. TBR collected the data. RD and PC analyzed the data and wrote the manuscript. All authors edited the manuscript.

## COMPETING INTERESTS

We have no competing interests.

## FUNDING

Supported by the National Science Foundation (NSF) IOS 1353085, the Smithsonian Migratory Bird Center, the Natural Sciences and Engineering Research Council of Canada (NSERC), and Carleton University.

## ACKNOWLEDGEMENTS

We thank Ben Vernasco, Camilo Alfonso, Brent Horton, Ignacio Moore, Brian Evans, David and Consuelo Romo, Kelly Swing, Diego Mosquera, Gabriela Vinueza, and the Tiputini Biodiversity Station of the Universidad San Francisco de Quito.

## Notes

### Competing Interest Statement

The authors have declared no competing interest.

### Summary of Updates

Slight revisions to clarify the text following one round of peer review

