## supplement for "Reciprocal social ties drive stability within a social network"

**Table S1. Descriptive statistics for one-directional and reciprocated partnerships.** Values from the observed data are shown first within each cell, followed by null estimates in italics (*null median ± SE*). The null permutations preserved each male’s spatiotemporal distribution of territory visits, but assumed that visits were independent of the presence or absence of other males. The undirected edge weight is based on the total interaction frequency between two partners in a given field season. For reciprocated partnerships, the mutual edge weight is the smaller of the two directed edge weights (i.e., the minimum contribution that occurred in both directions).

|  | 2015-16 | 2016-17 | 2017-18 |
| --- | --- | --- | --- |
| # of territory-holders tagged | 53 | 54 | 48 |
| **One-directional partnerships** |  |  |  |
| # Occurred | 44 *(47 ± 0.5)* | 57 *(69 ± 0.6)* | 34 *(29 ± 0.4)* |
| Average undirected edge weight | 6.5 *(2.0 ± 0.0)* | 11.1 *(1.6 ± 0.0)* | 11.1 *(1.6 ± 0.0)* |
| **Reciprocated partnerships** |  |  |  |
| # Occurred | 29 *(19 ± 0.2)* | 65 *(40 ± 0.3)* | 28 *(15 ± 0.2)* |
| Average undirected edge weight | 50.5 *(4.7 ± 0.0)* | 107.0 *(5.1 ± 0.0)* | 65.6 *(6.1 ± 0.1)* |
| # Occurred that had mutual edge weights > 5 | 14 *(0 ± 0)* | 26 *(0 ± 0.1)* | 11 *(0 ± 0.1)* |

**Table S2. Predictors of whether or not a partnership was reciprocated.** This binomial model evaluates the probability that a partnership was reciprocated vs. one-directional, as a binary response variable. The model accounts for field season, as well as three continuous predictors that were centred and scaled to have a mean of 0 and SD of 1 (edge weight, distance between the two territories, and recording duration). The random effect accounts for the identity of the dyad based on the combined IDs of the two participating males. To ensure that only partnerships that could have been reciprocated were considered, this analysis was limited to the 377 session partnerships that had an edge weight ≥2.

| **Fixed effects** | **Estimate** | **95% Confidence interval** | | **z** | **p-value** |
| --- | --- | --- | --- | --- | --- |
|  |  | **Lower** | **Upper** |  |  |
| Field season |  |  |  |  |  |
| 2015-16 | –0.93 | –1.54 | –0.33 |  |  |
| 2016-17 | –0.76 | –1.27 | –0.26 |  |  |
| 2017-18 | –1.21 | –1.92 | –0.49 |  |  |
| Edge weight (log) | 0.82 | 0.47 | 1.17 | 4.56 | < 0.0001 |
| Distance between territories | –2.97 | –4.17 | –1.77 | –4.85 | < 0.0001 |
| Recording duration | 0.58 | –0.003 | 1.16 | 1.95 | 0.05 |

**Table S3. Magnitude of individual donations in reciprocated partnerships.** This model evaluates whether the strength of social interactions a focal male donated to a particular partner was correlated with the amount he received from that same partner, within the same recording session. The response variable is ‘strength donated’ to a particular partner, relative to the focal male’s total interactions away from his own territory. The predictor ‘strength received’ (from that partner) is expressed relative to the focal male’s total interactions received on his own territory. The analysis also accounts for significant effects of field season, territory proximity (males donate more to closer neighbors), and the recording duration at the two respective territories; and it includes crossed random effects of both the focal and partner identities. Note that this analysis is limited to reciprocated session partnerships (n = 326 individual donations within 163 session-partnerships, among 72 males).

| **Fixed effect** | **Estimate** | **95% Confidence interval** | | **t** | **p** |
| --- | --- | --- | --- | --- | --- |
|  |  | **Lower** | **Upper** |  |  |
| Field season |  |  |  |  |  |
| 2015-16 | 0.44 | 0.30 | 0.58 |  |  |
| 2016-17 | 0.38 | 0.21 | 0.55 |  |  |
| 2017-18 | 0.52 | 0.37 | 0.67 |  |  |
| Strength received | 0.19 | 0.09 | 0.30 | 3.52 | 0.0005 |
| Distance between territories | –2.84 | –3.96 | –1.72 | –4.91 | < 0.0001 |
| Visits to donor by floater males (log pings) | 0.007 | –0.01 | 0.02 | 0.79 | 0.43 |
| Visits to recipient by floater males (log pings) | ­–0.01 | –0.03 | 0.002 | –1.70 | 0.09 |
| Recording duration | 0.09 | 0.03 | 0.14 | 3.13 | 0.002 |

**Table S4. Predictors of long-term partnership stability**. This model evaluates the stability of partnership edge weight through time. The response variable is a partnership’s edge weight in year *t + 1* (log-transformed). The predictors include field season, whether the partnership was reciprocated in year *t*, and its edge weight in year *t*. The random effect accounts for the identity of the dyad based on the combined IDs of the two participating males. This analysis was limited to n = 109 annual partnerships that had an edge weight ≥2, wherein both males were also tagged and monitored in the subsequent field season.

| **Fixed effect** | **Estimate** | **95% Confidence interval** | | **t** | **p-value** |
| --- | --- | --- | --- | --- | --- |
|  |  | **Lower** | **Upper** |  |  |
| Field season |  |  |  |  |  |
| 2015-16 | 0.98 | 0.19 | 1.77 |  |  |
| 2016-17 | –0.69 | –1.61 | 0.22 |  |  |
| Partnership type (year *t*, reciprocated vs. one-directional) | 0.91 | 0.18 | 1.63 | 2.46 | 0.02 |
| Distance between territories | –1.02 | –1.84 | –0.19 | –2.40 | 0.02 |
| Edge weight (year *t*, log) | 0.47 | 0.24 | 0.69 | 4.10 | < 0.0001 |
